# Actimetry in infant sleep research: an approach to facilitate comparability

**DOI:** 10.1101/494427

**Authors:** Sarah F. Schoch, Oskar G. Jenni, Malcolm Kohler, Salome Kurth

**Affiliations:** Pulmonary Clinic, University Hospital Zurich, Zurich, CH; Child Development Center, University Children’s Hospital Zurich, Zurich, CH; Children’s Research Center CRC, University Children’s Hospital Zurich, Zurich, CH

## Abstract

**Study Objectives:** Only standardized objective assessments reliably capture the large variability of sleep behavior in infancy, which is the most pronounced during the human lifespan. This is important for clinical as well as basic research. Actimetry is a cost-efficient method to objectively track infant sleep/wake behavior by assessing limb movements. Nevertheless, the standardization of actimetry-based sleep/wake measures is limited by two factors: the use of different computational approaches and the bias towards measuring only nighttime sleep - neglecting ∼20 % of sleep infants obtain during daytime.

**Methods:** We used actimetry in 50 infants for 10 continuous days at ages 3, 6 and 12 mo in a longitudinal approach. We analyzed the infants’ sleep/wake behaviors applying with two commonly used algorithms: Sadeh and Oakley/Respironics. We compared minute-by-minute agreement and Kappa between the two algorithms, as well as the algorithms with sleep/wake measures from a comprehensive 24-hour, parent-reported diary.

**Results:** Agreement between uncorrected algorithms was moderate (77 – 84%). By introducing a 6-step adjustment, we increased agreement between algorithms (96 – 97%) and with the diary. This decreased the difference between the two algorithms in *e.g. Total Sleep Duration* from 4.5 h to 0.2 h.

**Conclusions:** This computational pipeline enhances comparability between infant actimetry studies and the inclusion of parent-reported diaries allows the integration of daytime sleep. Objectively assessed infant sleep that is comparable across different studies supports the establishment of normative developmental trajectories and clinical cutoffs.

## INTRODUCTION

Studying the relationship between sleep in early life and later health and behavioral outcomes requires objective and reliable quantitative data. Current practice often relies on parent-reports to estimate infants sleep/wake behavior. However, subjective reports often disagree with objective sleep measures, *e.g*., misjudging sleep duration by >1h in young children ^1^. Wearables quantify sleep/wake states from arm or leg movement (actimetry) and allow cost-efficient sleep tracking in diverse environments and over long periods of time periods ^2^. Standardized procedures in actimetry studies will facilitate generalization of findings and cross-comparison between studies. Yet, we have to overcome two existing constraints: first, there are no standards for scoring sleep/wake from actimetry. The comparability of widely used analysis algorithms has never been investigated ^3^. Second, it is important to investigate both day- and night-time sleep in infants as sleep pressure and quality largely depend on the preceding history of day-/night-time sleep ^4^. Certain limitations (*e.g*., the underestimation of sleep due to external movements from carriage, stroller or bed-sharing, and the underestimation of wake when immobilized, *e.g*., baby sling, breastfeeding) have confined most infant actimetry assessments to nocturnal sleep, missing the ~20% of day-time sleep ^5^.

We compute the sleep- or wake-bias of two common approaches and present adaptations to streamline sleep/wake scoring and quantify infant daytime sleep by integrating detailed 24-h diary information into actimetry analysis ^6^. Applying this approach will facilitate the comparability across studies.

## METHODS

### Participants

50 healthy infants (17 female) were longitudinally assessed with ankle actimetry at age 3 mo (*i.e*., 2.46 – 3.38 mo at assessment start), 6 mo (5.42 – 6.18 mo) and 12 mo (11.47 – 12.16 mo). The presence of medical conditions and travelling across time zones with >1 hour difference in the 4 weeks prior to assessment served as exclusion criteria. Ethical approval was obtained from the *Zurich ethics committee (2016-00730)* and study procedures were consistent with the declaration of Helsinki. Written parental consent was obtained before enrollment.

### Experimental design

Data was collected at each assessment time point for a duration of 10 days (5-16 d) through ankle actimetry and a 24h sleep-wake diary. GENEactiv accelerometers (Activinsights Ltd, Kimbolton, UK; 43×40×13 mm, MEMS sensor, 16 g, 30 Hz frequency; sensitive for +/- 8 g range at 3.9 mg resolution) were attached on the left ankle with a modified sock or a Tyvek paper strap. Parents were instructed to only remove the actimeter for bathing and to document its removal in the 24-h diary. The sleep diary was adapted from Werner, Molinari ^1^, with parents reporting in 15 min intervals: sleep (including external movement, *e.g*., sleeping in the parents arms, stroller *etc*.), wake, feeding, and crying. Parents reported bed times (putting infant to bed in the evening and getting up in the morning), naps and marked particular periods of uncertainty (*e.g.* feeding periods during nighttime). They were instructed to fill out the diary throughout the day. During the assessment, the Brief Infant Sleep Questionnaire (BISQ) was completed ^7^. Families received small gifts for the infant (*i.e.* bottles, baby food) for participation.

### Actimetry processing

Actimetric data was extracted as binary files using GENEactiv PC Software (Version 3.1), imported into Matlab (R2016b), and converted to activity counts ^8^, including a 3-11 Hz bandpass filter and signal compression to 15 s bins. Acceleration data from the three axes was combined using a sum of squares. Signal was compressed to one data point per minute by data summation. To identify infant sleep and wake periods, several adjustments were introduced to existing algorithms (Sadeh, Acebo ^9^ and Oakley ^10^, Figure 1A). First, the threshold value was changed. In the Oakley algorithm generally a threshold of 20, 40 or 80 is used, this was replaced with mean activity of the full recording*0.88 (similar to the auto-threshold in Respironic devices). In the Sadeh algorithm 100 is used to distinguish between a low and high activity epoch, this was also replaced by mean activity of the full recording*0.88. As the original algorithms revealed strong bias to either sleep or wake, in a second step we added (Sadeh)/subtracted (Okaley) a factor based on mean activity of each recording ^11^ (Figure 1B). Third, data was adapted and rescored with information from the 24-h diary. Time periods when the actimeter was not worn were scored with reports from the 24-h diary (Figure 1C). Mis-scoring of sleep due to short periods of inactivity during wake was rescored according to strict criteria (short periods of sleep surrounded by certain periods of wake, and the first 1-4 minutes of sleep are rescored wake depending on the length of the wake period before, Figure 1D) by Webster, Kripke ^12^. Diary-documented sleep with external movement was re-scored as sleep (Figure 1E). Next, wake periods <5 min which bilaterally bordered scored sleep where assigned to sleep (Smoothing, Figure 1F).

**Figure 1.**
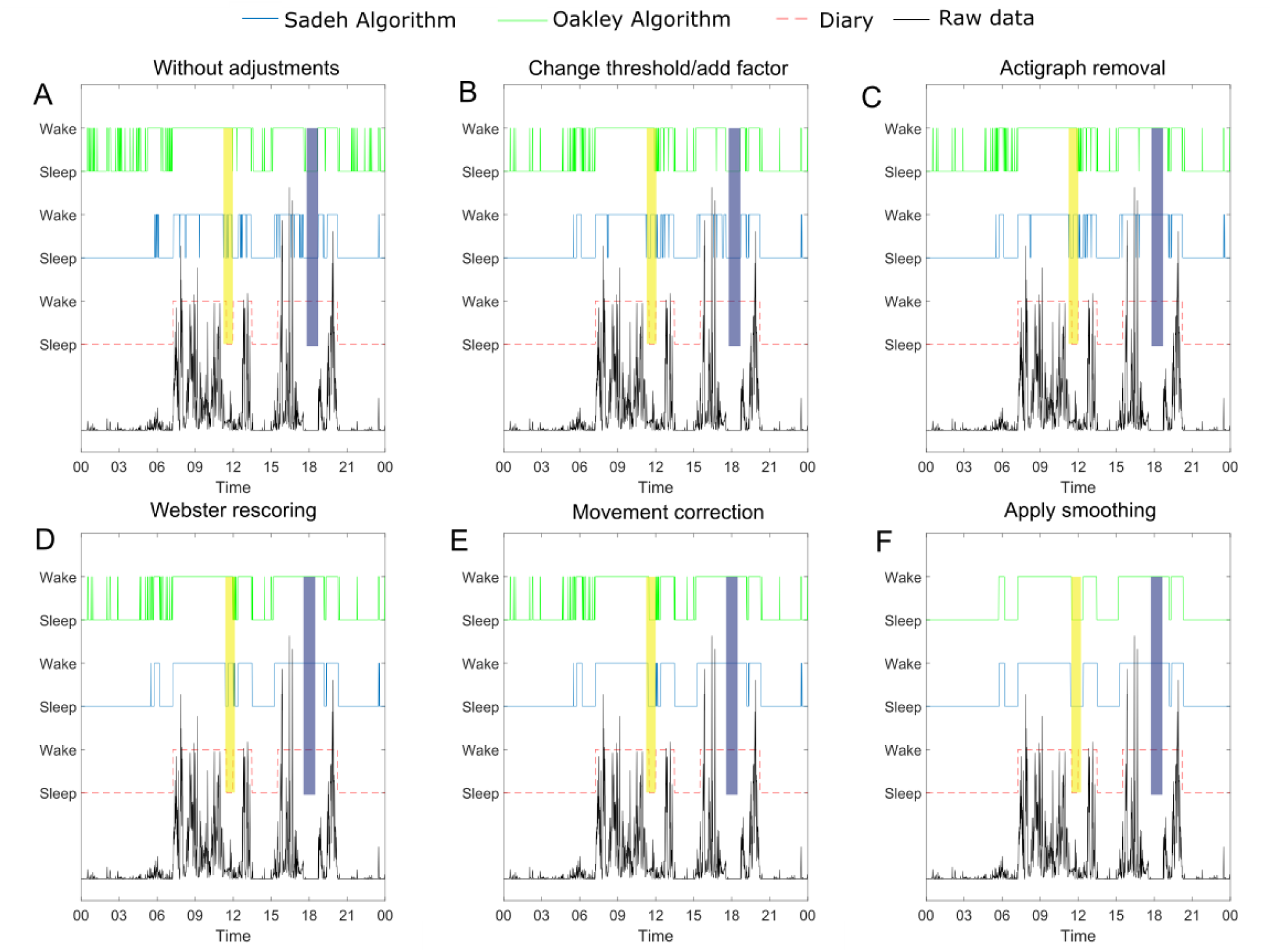
Stepwise processing adjustments. Typical 24-hour actimetric profile from a representative participant (age 12 mo). Raw data (black) and the scorings from 24-h diary (red), Sadeh (blue) and Oakley (green) are presented. Wake is shown on top and sleep at the bottom of each scoring item. Stepwise adjustments are presented in order of processing: A) Raw data without adjustments; B) Altered threshold and added factor reducing wake/sleep bias; C) Rescoring of actimeter removal with 24-h diary information; D) Rescoring by Webster; E) Rescoring of sleep with external movements; and F) Smoothing of short wake periods (< 5 min) during sleep. Yellow shading indicates periods of sleep with reported external movement. Blue shading illustrates periods with actimeter removal.

In order to reduce error caused by external factors, 24-hour days were excluded for the calculation of sleep variables if 1) the actimeter was removed for > 3 h (in 22.2 % of all data including first and last days of the assessment, and interruptions due to sickness), 2) the infant was sick but the overall assessment was continued (4.2 %), or 3) the assessment took place during the switch to/from daylight savings (0.2%). These criteria resulted in the following data included in final analysis: mean assessment duration of 8.6 ± 1.65 d at age 3 mo (whereby 3 d was the minimum assessment duration and 13 d the maximum), correspondingly 8.0 ± 1.95 d included at 6 mo (2 – 11d) and 7.9 ± 1.71 d included at 12 mo (3 – 10d).

From the resulting matrix containing a minute-by-minute scoring of either sleep or wake, sleep variables of interest were computed: *Total Sleep Duration, Day-to-Day Sleep Variability, % Night Sleep, Fragmentation*. *Total Sleep Duration (h)* sums the time scored as sleep within 24 h (starting at clock time 0:01). *Day-to-Day Sleep Variability (h)* is the standard deviation of the *Total Sleep Duration* across all assessment days. *% Night Sleep* indicates the relative proportion of night-time sleep (*i.e*., within clock time 19:00 – 07:00) as a percentage of *Total Sleep Duration*. *Fragmentation* (awakenings/h) calculates the number of awakenings per hour during nighttime sleep (based on individual infant bedtimes reported by parents). Awakenings were scored separate when divided by at least 10 min of sleep. BISQ total sleep duration was calculated by adding reported day and night sleep duration (rounded to 15 min; mean was used when time range was reported). N = 9 BISQ assessments were excluded due to incomplete data.

### Statistical analysis

We used R (version 3.3.2) and R Studio (version 1.0.136) for statistical analyses. Linear mixed-effect models were estimated using restricted maximum likelihood to analyze changes resulting from adjustments using the R-packages *lmer* ^13^ and *lmertest* ^14^. The covariate assessment time point was included as a logarithmic function of age (log(age)). We chose this logarithmic function to account for the flattening of effects with age (larger effects between 3 and 6 months than between 6 and 12 months). All models included effects of adjustment, infant age and their interaction. To compare whether random effects of time point and adjustments improve model fit we compared one model combining both random effects with two separate models containing random effects of either time point or adjustment. The random effects where only included in the final model if it significantly improved the model fit with most weight given to the Bayes Information Criterion (BIC, Tables 1-5, selected model highlighted in bold). We calculated agreement between two measures as % of 1-min periods scoring the same state (*i.e*., sleep or wake) and additionally using Cohen’s Kappa ^15^. Bias was calculated as the difference (min) where one algorithm scored sleep and the other wake. We used Bland Altman statistics to investigate whether the algorithms calculated similar estimates for sleep variables (package *BlandAltmanLeh*). A two-sided significance level of P < 0.05 was used.

## RESULTS

### Agreement between algorithms

We compared the agreement between the algorithms with and without adjustments. Without adjustment, algorithms show moderate agreement in scoring sleep or wake (77 – 84%, κ = 0.50 – 0.68, Table 6). Agreement was significantly improved by introducing the 6-step adjustment (96-97%, κ = 0.91 – 0.95, t_(274.63)_ = 23.35, P < 0.0001, Table 1). The largest disagreement was observed in actimetry data from infants age 3 mo (t_(247)_ = 14.44, P < 0.0001). The largest improvement in agreement occurred at age 3 mo (interaction age * improvements, t_(247)_ = −7.63, P < 0.0001). The improved agreement mainly results from threshold adaptation and adding the factor against bias (~5%) as well as smoothing (~2%).

**Table 6.** Agreement rates with and without adjustment steps.

| Age | No adjustments | Change threshold | Add factor | Actigraph removal | Rescoring Webster | External movements | Smoothing |
| --- | --- | --- | --- | --- | --- | --- | --- |
| Sadeh - Oakley |  |  |  |  |  |  |  |
| 3 months | 77.36 ± 3.97 | 83.21 ± 3.75 | 91.56 ± 1.59 | 91.64 ± 1.59 | 92.96 ± 1.34 | 93.56 ± 1.38 | 95.76 ± 1.00 |
| 6 months | 83.65 ± 3.67 | 87.65 ± 3.68 | 93.14 ± 1.34 | 93.23 ± 1.33 | 93.43 ± 1.13 | 94.89 ± 1.16 | 96.79 ± 0.80 |
| 12 months | 83.97 ± 3.07 | 87.99 ± 3.12 | 93.66 ± 1.32 | 93.78 ± 1.32 | 94.66 ± 1.34 | 94.88 ± 1.36 | 97.43 ± 0.83 |
| Sadeh - Diary |  |  |  |  |  |  |  |
| 3 months | 76.39 ± 6.05 | 75.25 ± 6.11 | 79.55 ± 5.19 | 80.22 ± 5.16 | 81.65 ± 5.17 | 86.22 ± 5.17 | 86.36 ± 5.20 |
| 6 months | 82.26 ± 4.93 | 81.18 ± 4.92 | 84.42 ± 3.59 | 85.67 ± 3.60 | 86.86 ± 3.59 | 89.60 ± 3.21 | 89.69 ± 3.26 |
| 12 months | 85.40 ± 6.23 | 84.30 ± 6.12 | 88.19 ± 5.11 | 90.21 ± 2.87 | 91.55 ± 2.66 | 93.07 ± 2.42 | 93.23 ± 2.44 |
| Oakley - Diary |  |  |  |  |  |  |  |
| 3 months | 75.14 ± 4.57 | 77.17 ± 4.83 | 77.22 ± 4.86 | 77.91 ± 4.81 | 79.24 ± 7.40 | 84.05 ± 4.95 | 85.73 ± 5.08 |
| 6 months | 80.77 ± 3.34 | 82.77 ± 3.53 | 82.77 ± 3.55 | 84.00 ± 3.56 | 84.76 ± 3.41 | 87.85 ± 3.21 | 89.45 ± 3.25 |
| 12 months | 84.15 ± 5.04 | 86.34 ± 5.11 | 86.36 ± 5.13 | 88.38 ± 2.83 | 89.08 ± 2.86 | 90.76 ± 2.72 | 92.99 ± 2.50 |

*Agreement rates as % agreement (averaged over participants over measurement days). Agreements are shown between the two algorithms and between each algorithm and sleep diaries filled out by the parents.*
*Note: Means ± SD is shown.*

### Agreement between algorithms and 24-h diary

We compared the scorings of both algorithms, with and without adjustments, with the parental reported 24-h diary. Both algorithms showed medium agreement with the 24-h diary without adjustments (75 – 85%, Sadeh vs Diary κ = 0.51 – 0.70, Oakley vs Diary κ = 0.5 – 0.68, Table 6). Adjustments increased agreement to up to 93% (86 - 93%, Sadeh vs Diary κ = 0.72 – 0.86, Oakley vs Diary κ = 0.71 – 0.86, t_(494.42)_ = 13.93, P < 0.0001, Table 2). Lower agreement was seen for 3 mo olds compared to 6 and 12 mo olds (t_(495)_ = 12.19, P < 0.0001). An interaction between age and improvement (t_(495)_ = −3.31, P = 0.001), indicates greater improvements in the youngest age group. There was no significant effect of algorithm (Sadeh vs Oakley t_(1,495)_ = 1.90, P = 0.06) and no interaction of type of algorithm and age (t_(495)_ = −0.32, P = 0.75). A small interaction was observed between algorithm and amount of improvements, with the Oakley algorithm showing increased improvements due to the adjustments (t_(495)_ = −2.02, P = 0.04). At 3 mo, adjusting for movement during sleep greatly improved the agreement (~4.5%), which was less pronounced for 6 and 12 mo respectively (~1.5 – 3 %). The opposite was seen for adjustments for actimeter removal, which occurred less at 3 mo (0.68%) than at 6 and 12 mo (~1.25 – 2%).

### Bias towards sleep or wake

Each algorithm had a scoring bias for a specific state: 200-300 mins per day were scored as sleep by the Sadeh algorithm and wake by the Oakley algorithm (Figure 2). This bias was significantly reduced by the adjustments (t_(282.49)_ = −27.34, P < 0.0001, Table 3). Particularly 3 months-olds’ scorings showed increased bias in comparison with the older infants’ scorings (t_(247)_ = −13.35, P < 0.0001), but bias decreased most through our adjustments at that age (t_(1,247)_ = 11.45, P < 0.0001). Similar bias was observed when compared to the 24-h diary: the Sadeh algorithm scored more sleep than reported in the 24-h diary. This bias decreased by the adjustments (t_(198)_ = −6.38, P < 0.0001). The bias was stronger with lower age (t_(100.58)_ = −2.79, P = 0.006) but showed no interaction with age (t_(198)_ = 1.50, P = 0.13). The Oakley algorithm scored more wake compared to the sleep 24-h diary. This bias was significantly reduced by our adjustments (t_(81.81)_ = 11.68, P < 0.0001). There was an age effect (t_(90.08)_ = 3.50, P = 0.0007), with the largest improvements at 3 mo (t_(198)_ = −5.75, P < 0.0001).

**Figure 2.**
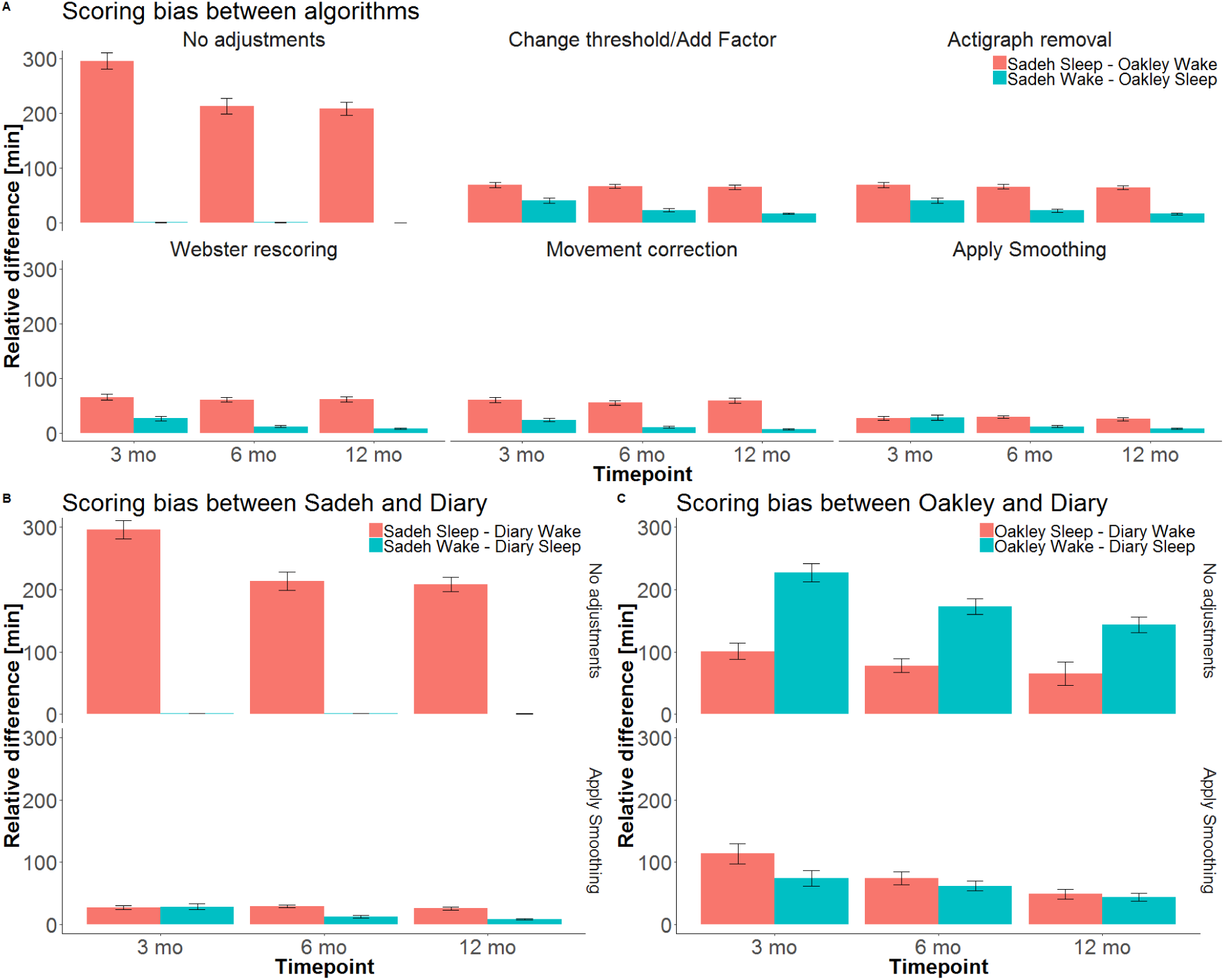
Sleep/wake bias of scoring algorithms and 24-hour-diary reported by parents. Scoring bias shows disagreement of scoring as sleep or wake (sum of minutes within 24h, error bars represent 95% confidence interval). A) Scoring bias is depicted without adjustments and for each adjustment step. B) Scoring bias of the Sadeh algorithm and compared to the 24-h diary is shown without adjustments and including all adjustments. C) Bias of the Oakley algorithm compared to the diary is shown without adjustments and including all adjustments.

### Sleep/Wake behavior estimation

To estimate differences in the sleep parameters, we calculated Bland Altman statistics of each parameter without and with adjustments (Table 7 and figures 3-6). Without adjustments, there was a bias in the variables *Sleep Duration, % Night Sleep* and *Fragmentation*, as shown by data points instead of being centered around 0 (no bias) they were centered around *e.g.* 4.2 h for *Sleep Duration* (Figure 3). Bias for each age group is shown in Table 2, e.g. 5.45 h at age 3 months (previously this approach was used with a bias definition exceeding ± 0.5 h ^1^). This bias was reduced by our adjustments, to *e.g.* −0.01 h in *Sleep Duration* at 3 months. The only variable showing low bias (mean < 0.5 h) already without adjustments was *Day-to-Day sleep variability*. Taken together we show that infant actimetry-based detection of sleep/wake variables can be improved by 6-steps of adjustments.

**Table 7.** Differences in sleep parameter estimates with and without adjustments estimated by Bland Altman scores.

| Age | Sleep Duration [h] |  | Day-to-day Sleep Variability [h] |  | % Night Sleep |  | Fragmentation [/h] |  |
| --- | --- | --- | --- | --- | --- | --- | --- | --- |
|  | No adjustments | After adjustments | No adjustments | After adjustments | No adjustments | After adjustments | No adjustments | After adjustments |
| 3 months | 5.45 ± 1.90 | -0.01 ± 0.90 | -0.09 ± 0.70 | 0.01 ± 0.22 | -6.99 ± 5.74 | 0.11 ± 1.52 | -0.65 ± 0.37 | -0.04 ± 0.09 |
| 6 months | 3.96 ± 1.69 | 0.33 ± 0.41 | -0.06 ± 0.81 | 0.00 ± 0.22 | -6.64 ± 6.33 | -0.40 ± 1.08 | -0.78 ± 0.35 | -0.06 ± 0.12 |
| 12 months | 4.08 ± 1.46 | 0.32 ± 0.37 | -0.09 ± 0.55 | -0.09 ± 0.20 | -4.74 ± 5.60 | 3.35 ± 1.34 | -0.92 ± 0.32 | -0.17 ± 0.08 |

Note: Means ± Critical Difference is shown. Means show general bias of one measure over the other. Critical difference show 95% differences in estimates.

**Figure 3.**
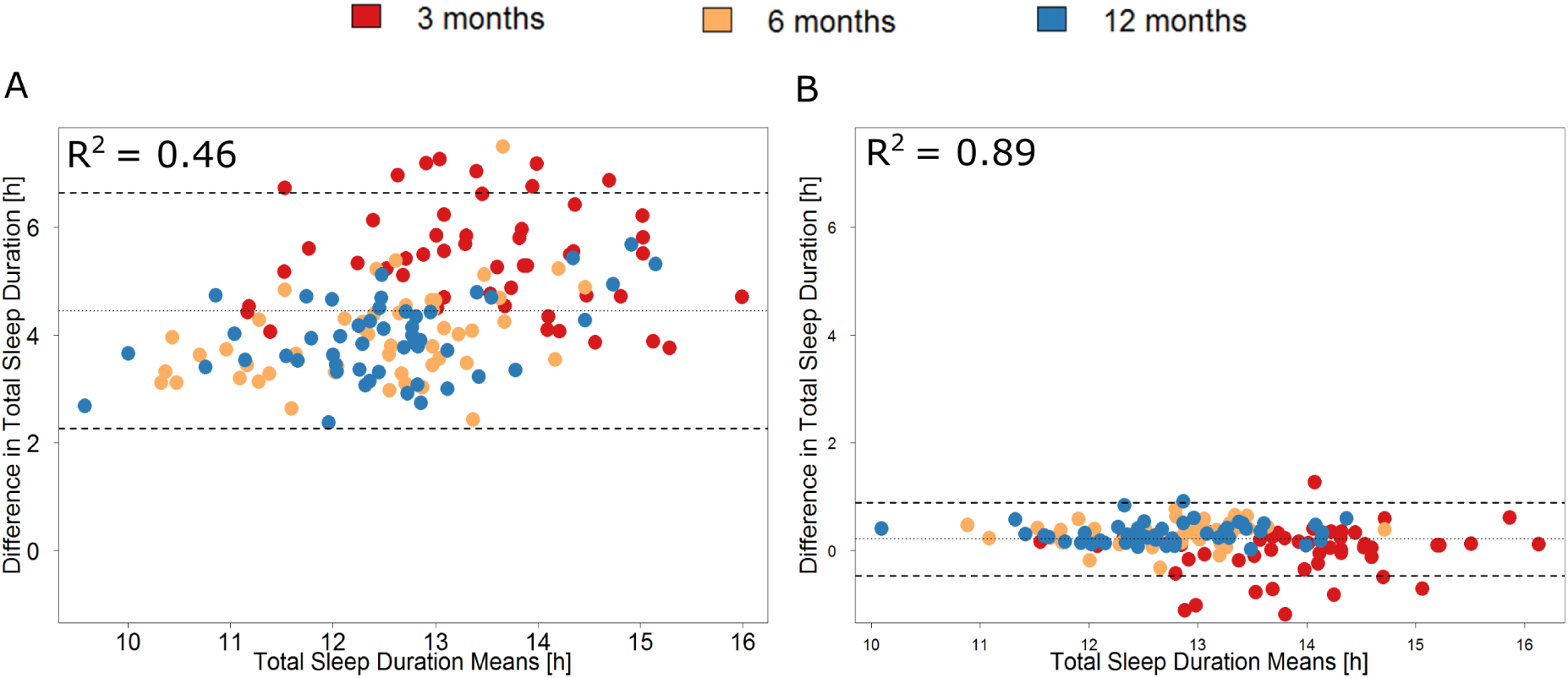
Bland-Altman plots of *Total Sleep Duration* estimates from Sadeh and Oakley algorithm. Each infant is represented by three dots indicating the age group by color. A) Without adjustments, the difference in *Total Sleep Duration* is > 4 h with a critical difference of 2.18 h, indicating that without adjustments *Total Sleep Duration* estimates from the Sadeh algorithm are 2 to 7 hours above estimates from the Oakley algorithm. B) With 6-step adjustments, the difference in *Total Sleep Duration* is lowered to ~ 0 h with a critical difference of 0.68 h.

**Figure 4.**
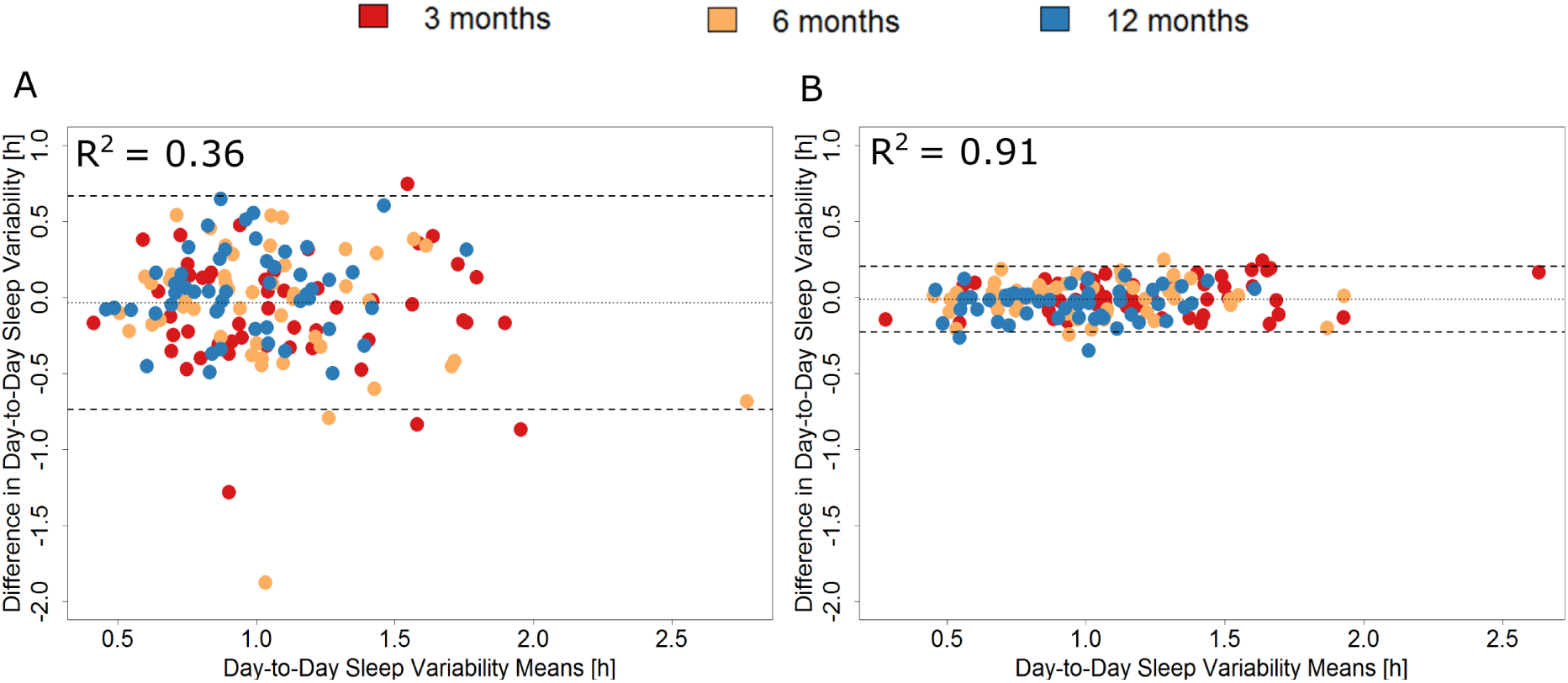
Bland-Altman plots showing difference in *Day-to-Day Sleep Variability* between scoring based on Sadeh and Oakley algorithms. Each infant is represented by three dots indicating the age group by color. A) Without adjustments, the difference in *Day-to-Day Sleep Variability* is ~0 h with a critical difference of 0.7 h. B) With 6-step adjustments, the difference in *Day-to-Day Sleep Variability* estimate is ~ 0 h with a critical difference of 0.22 h.

**Figure 5.**
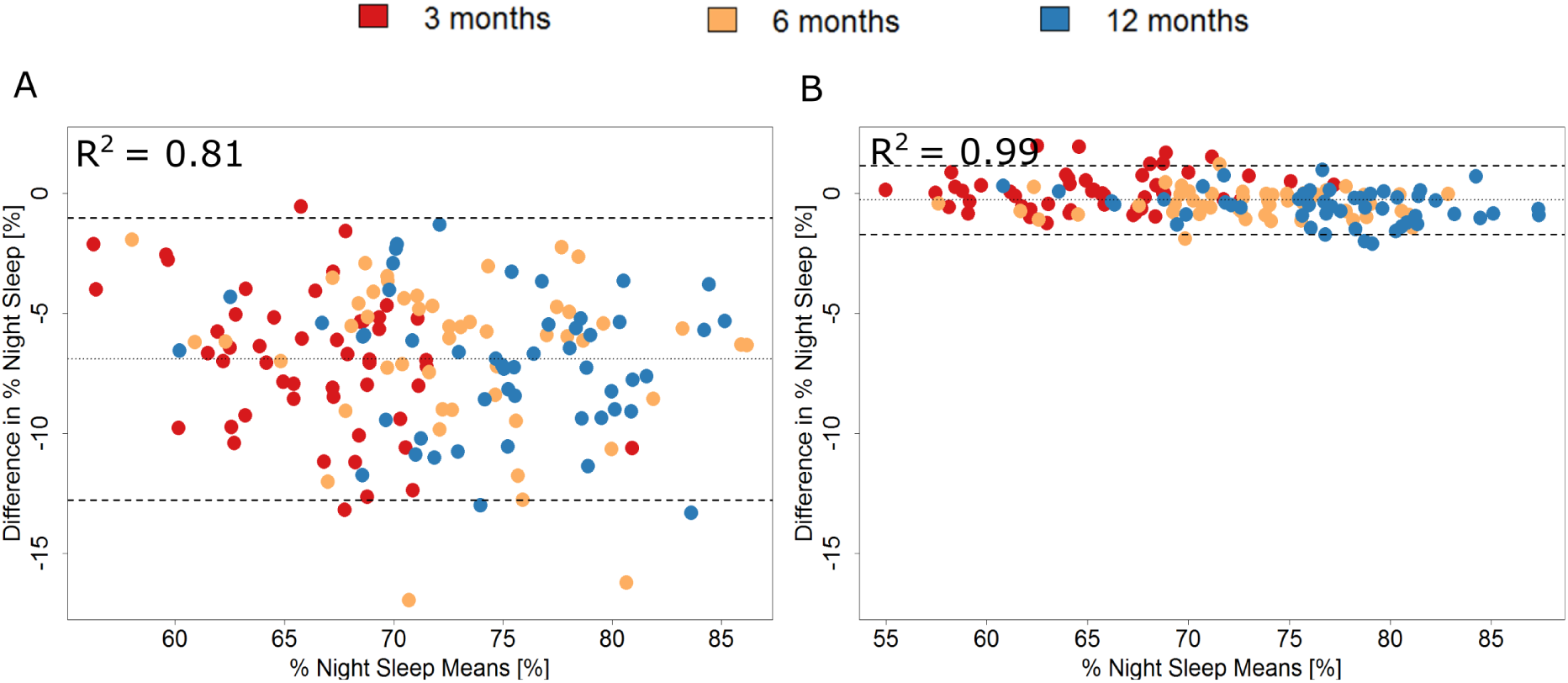
Bland-Altman plots showing difference in *% Night Sleep* between scoring based on Sadeh and Oakley algorithm. Each infant is represented by three dots indicating the age group by color. A) Without adjustments the mean difference in *% Night Sleep* estimate is ~7% with a critical difference of 5.87%. B) With 6-step adjustments the mean difference in *% Night Sleep* estimate is ~ 0% with a critical difference of 1.43%.

**Figure 6.**
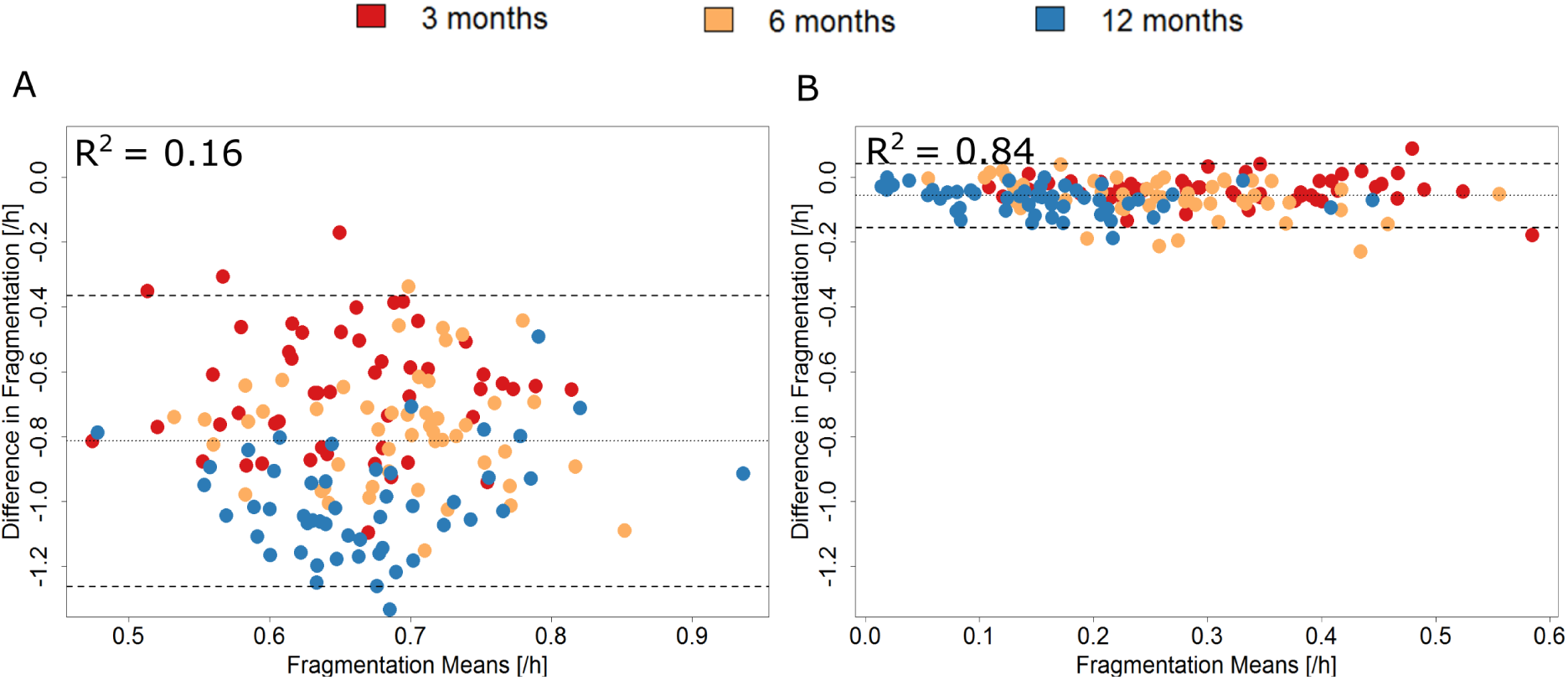
Bland-Altman plots showing difference in *Fragmentation* between scoring based on Sadeh and Oakley algorithm. Each infant is represented by three dots indicating the age group by color. A) Without adjustments the mean difference in *Fragmentation* estimate is ~0.8% with a critical difference of 0.45. B) With 6-step adjustments the mean difference in *Fragmentation* estimate is ~ 0 with a critical difference of 0.1%.

### Comparison to questionnaire data

We found a generally large deviation from parental questionnaire data compared to actimetry data in *Total Sleep Duration*, as indicated by a critical difference of 3.19 h. This includes both, under- and overestimating of the objective estimates by parent’s estimates (95%; see supplementary Figures 1 and 2). For example, parents reporting their infants’ sleep duration to be 13.5 h, revealed objectively measured infant sleep duration between 11.86 h and 14.6 h. However, there was no systematic bias (i.e. either under- or overestimating) *Total Sleep Duration* of their infants.

## DISCUSSION

Only standardized objective assessments reliably capture the large variability of sleep behavior in infancy, which is the most pronounced during the human lifespan ^5^. We optimized sleep quantification from actimetry in infants by applying a set of adjustments that overcomes discrepancies in sleep measures between existing scoring algorithms ^9^, ^10^. The use of 24-h diaries minimizes signal miscomputation through external factors and improves the analysis of daytime sleep. These methods will help to extend reference values based on parental reports ^5^ or meta-analysis based on different devices ^16^.

Adjustments reduce disagreement between algorithms from 16-22% to 3-4%. In addition, the inherent bias of the two algorithms (Sadeh algorithm towards sleep; Oakley algorithm towards wake) was significantly reduced by the adjustments. Such standardization is of great importance for computation of sleep variables. For example, without adjustments, *Total Sleep Duration* deviates up to 7 h depending on the algorithm used, with higher sleep duration estimates when using the Sadeh algorithm compared to the Oakley algorithm. After adjustments, these estimates vary less than 1 h. Importantly this also increased the correlation, meaning that the infants which overall showed the highest sleep duration as calculated from one algorithm also are estimated to have a high sleep duration with the other algorithm. Similar effects were seen for parameters such as *% Night Sleep* and *Fragmentation.* Only *Day-to-Day Sleep Variability* showed no bias without adjustments, but even for this parameter correlation could be improved drastically.

We also identified age-specific effects that affect actimetry outcomes. Scoring agreement generally increases when infants with age. We hypothesize that this is primarily due to increased motor activity during wake as part of motor development. Additionally, external movement during sleep in very young infants can lead to mis-scoring of up to 1 h. This was corrected by introducing our adjustments. Furthermore, with increasing age, removal time of the actimeters increased (*e.g.* removal by child or other infants, longer periods of bathing/water activities), which led to mis-scoring of up to 30 minutes. Completing the 24-hour diary remains important for the reliable detection and correction of such incidents.

Moreover, our approach integrates the ~20% of infant sleep occurring during daytime. Daytime naps are often missed in traditional analyses, but they reflect the important build-up of sleep pressure and the neurophysiological capacity of children to increase consolidated waking bouts ^17^. Our approach circumvents these difficulties by integrating complementary information from a 24-hour sleep diary. Although our semi-automated integration requires time investment of study participants and researchers, it greatly improves data reliability and allows comparison across studies. We suggest to integrate digital diaries (*i.e.* sleep tracking apps) linked to actimetry input for future studies. Parents should be given the opportunity to confirm sleep periods or reject faulty ones electronically. Additional computational corrections can be introduced to i) distinguish between movements of the infant vs. external movements, or ii) automatically detect periods where the actimeter is not worn. This requires the integration of new sensors such as heart rate or skin temperature. Such sensors could also distinguish quiet wakefulness from sleep, which cannot be achieved with acceleration only.

In conclusion, we present adjustments to standardize actimetric sleep/wake scoring for nighttime and daytime sleep. Applying these adjustments increases the reliability of measured infant sleep variables.

## Supporting information

## Acknowledgments

This work was supported by the Clinical Research Priority Program Sleep and Health of the University of Zurich (to S.K.) and Swiss National Science Foundation (P0ZHP1-178697 to S.F.S).

Financial Disclosure: none.

Non-financial Disclosure: none.

