## Supplementary material for "Actimetry in infant sleep research: an approach to facilitate comparability"

**Table 1**. Parameters for the models examining agreement between Sadeh and Oakley algorithm. Model 2 that only includes the random effects of the adjustments shows the best model fit.

|  | Model 1 | **Model 2** | Model 3 |
| --- | --- | --- | --- |
| *Fixed effects* | b ± SE [95% CI] | **b ± SE [95% CI]** | b ± SE [95% CI] |
| Intercept | 73.12 ± 0.79 [71.58; 74.66] | **73.12 ± 0.70 [71.76; 74.49]** | 73.12 ± 0.76 [71.63; 74.61] |
| Adjustments | 21.39 ± 0.88 [19.66; 23.11] | **21.39 ± 0.92 [19.59; 23.18]** | 21.39 ± 0.97 [19.48; 23.29] |
| Age | 4.77 ± 0.35 [4.08; 5.46] | **4.77 ± 0.33 [4.12; 5.41]** | 4.77 ± 0.37 [4.03; 5.51] |
| Adjustements *Age | -3.56 ± 0.45 [-4.44; -2.67] | **-3.56 ± 0.47 [-4.48; -2.65]** | - 3.56 ± 0.52 [-4.58; -2.55] |
| *Random effects* | b | **b** | b |
| Intercept | 13.43 | **5.02** | 5.16 |
| Adjustments | 3.59 | **3.50** |  |
| Age | 1.22 |  | 0.42 |
| AIC | 1428.3 | **1428.6** | 1462.9 |
| BIC | 1469.0 | **1458.2** | 1492.5 |
| Pseudo R^2^ (marginal; conditional) |  | **0.89; 0.92** | Model 2 |

Note: *AIC = Akaike Information Criterion, BIC = Bayes Information Criterion, SE = Standard Error*

**Table 2**. Parameters for the models examining agreement between the algorithms and the diary. Model 1 that includes both random effects shows the best model fit.

|  | **Model 1** | Model 2 | Model 3 |
| --- | --- | --- | --- |
| *Fixed effects* | **b ± SE [95% CI]** | b ± SE [95% CI] | b ± SE [95% CI] |
| Intercept | **68.25 ± 1.20 [65.90; 70.60]** | 68.25 ± 0.94 [66.41; 70.10] | 68.25 ± 1.21 [65.88; 70.62] |
| Adjustments | **11.89 ± 0.85 [10.21; 13.56]** | 11.89 ± 0.97 [9.98; 13.79] | 11.89 ± 0.86 [10.20; 13.57] |
| Age | **6.57 ± 0.54 [5.51; 7.62]** | 6.57 ± 0.43 [5.73; 7.40] | 6.57 ± 0.54 [5.50; 7.63] |
| Algorithm (Sadeh vs Oakley) | **1.58** ± **0.83 [-0.05; 3.21]** | 1.58 ± 0.97 [-0.31; 3.47] | 1.58 ± 0.86 [-0.10; 3.26] |
| Adjustments * age | **-1.40 ± 0.42 [-2.3; -0.57]** | -1.40 ± 0.49 [-2.37; -0.44] | -1.40 ± 0.44 [-2.26; -0.56] |
| Age * Algorithm | **-0.14 ± 0.42 [-0.97; 0.69]** | -0.14 ± 0.49 [-1.10; 0.83] | -0.14 ± 0.44 [-1.00; 0.71] |
| Adjustments * Algorithm | **-0.97 ± 0.48 [-1.91; -0.02]** | -0.97 ± 0.56 [-2.06; 0.12] | -0.97 ± 0.49 [-1.94; 0.002] |
| *Random effects* | **b** | b | b |
| Intercept | **47.60** | 11.24 | 47.00 |
| Adjustments | **1.90** | 0.85 |  |
| Age | **7.80** |  | 7.62 |
| AIC | **3218.4** | 3304.9 | 3237.5 |
| BIC | **3280.0** | 3353.3 | 3285.8 |
| Pseudo R^2^ (marginal; conditional) | **060; 0.83** |  |  |

Note: *AIC = Akaike Information Criterion, BIC = Bayes Information Criterion, SE = Standard Error*

**Table 3**. Parameters for the models examining bias between the algorithms. Model 2 that includes only adjustments as random effects shows the best model fit.

|  | Model 1 | **Model 2** | Model 2 |
| --- | --- | --- | --- |
| *Fixed effects* | b ± SE [95% CI] | **b ± SE [95% CI]** | b ± SE [95% CI] |
| Intercept | 351.20 ± 9.88 [331.76; 370.48] | **351.20 ± 9.74 [332.02; 370.22]** | 351.20 ± 10.41 [330.72; 371.52] |
| Adjustments | -363.89 ± 13.31 [- 389.96; -337.81] | **-363.89 ± 13.31 [- 389.97; -337.81]** | -363.89 ± 14.69 [- 392.68; -335.1] |
| Age | -62.82 ± 4.71 [-72.03; -53.59] | **-62.82 ± 4.70 [-72.03; -53.60]** | -62.82 ± 5.53 [-73.65; -51.98] |
| Adjustments * age | 76.15 ± 6.65 [63.12; 89.19] | **76.15 ± 6.65 [63.11; 89.19]** | 76.15 ± 7.82 [60.83; 91.47] |
| *Random effects* | b | **b** | b |
| Intercept | 973.98 | **839.7** | 19.55 |
| Adjustments | 1042.97 | **1041.6** |  |
| Age | 1.55 |  | 0.02 |
| AIC | 3021.1 | **3015.2** | 3054.3 |
| BIC | 3061.9 | **3044.9** | 3084.0 |
| Pseudo R^2^ (marginal; conditional) |  | **0.90; 0.93** |  |

Note: *AIC = Akaike Information Criterion, BIC = Bayes Information Criterion, SE = Standard Error*

**Table 4**. Parameters for the models examining bias between the Sadeh algorithm and the diary. Model 3 that includes only age as random effects shows the best model fit.

|  | Model 1 | Model 2 | **Model 3** |
| --- | --- | --- | --- |
| *Fixed effects* | b ± SE [95% CI] | b ± SE [95% CI] | **b ± SE [95% CI]** |
| Intercept | 190.19 ± 21.16 [148.72; 231.66] | 190.19 ± 17.99 [154.93; 225.45] | **190.19 ± 20.76 [149.49; 230.89]** |
| Adjustments | -140.05 ± 21.21 [- 181.62; -98.48] | -140.05 ± 23.81 [- 186.71; -93.39] | **-140.05 ± 21.95 [- 183.06; -97.04]** |
| Age | -28.79 ± 10.16 [-48.70; -8.88] | -28.79 ± 8.85 [-46.13; -11.45] | **-28.79 ±10.32 [-49.01; -8.57]** |
| Adjustments * age | 17.54 ± 11.09 [-4.19; 39.27] | 17.54 ± 12.51 [-6.98; 42.07] | **17.54 ± 11.68 [-5.35; 40.44]** |
| *Random effects* | b | b | **b** |
| Intercept | 11532.5 | 2364.1 | **9518** |
| Adjustments | 788.3 | 694.3 |  |
| Age | 2086.3 |  | **1910** |
| AIC | 3379.3 | 3392.5 | **3387.8** |
| BIC | 3420.0 | 3422.1 | **3417.4** |
| Pseudo R^2^ (marginal; conditional) |  |  | 0.37; 0.61 |

Note: *AIC = Akaike Information Criterion, BIC = Bayes Information Criterion, SE = Standard Error*

**Table 5**. Parameters for the models examining bias between the Oakley algorithm and the diary. Model 3 that includes only age as random effects shows the best model fit.

|  | Model 1 | Model 2 | **Model 3** |
| --- | --- | --- | --- |
| *Fixed effects* | b ± SE [95% CI] | b ± SE [95% CI] | **b ± SE [95% CI]** |
| Intercept | -160.84 ± 20.01 [-200.06; -121.62] | -160.84 ± 16.71 [-193.59; -128.08] | **-160.84 ± 19.77 [-199.58; -122.10]** |
| Adjustments | 223.74 ± 18.51 [187.46; 260.02] | 223.74 ± 21.69 [181.23; 266.25] | **223.74 ± 19.16 [186.19; 261.29]** |
| Age | 34.02 ± 9.59 [15.22; -52.81] | 34.02 ± 8.07 [18.20; -49.84] | **34.02 ± 9.72 [14.97; 53.06]** |
| Adjustments * age | -58.59 ± 9.67 [-77.54; -39.64] | -58.59 ± 11.41 [-80.96; -36.22] | **-58.59 ± 10.20 [-78.58; -38.61]** |
| *Random effects* | b | b | **b** |
| Intercept | 11769.3 | 2466 | **10354** |
| Adjustments | 630.2 | 521 |  |
| Age | 2260.9 |  | **2122** |
| AIC | 3323.8 | 3323.8 | **3333.9** |
| BIC | 3365.6 | 3365.6 | **3363.6** |
| Pseudo R^2^ (marginal; conditional) |  |  | **0.44;0.71** |

Note: *AIC = Akaike Information Criterion, BIC = Bayes Information Criterion, SE = Standard Error*
