## Supplementary material for "Actimetry in infant sleep research: an approach to facilitate comparability"

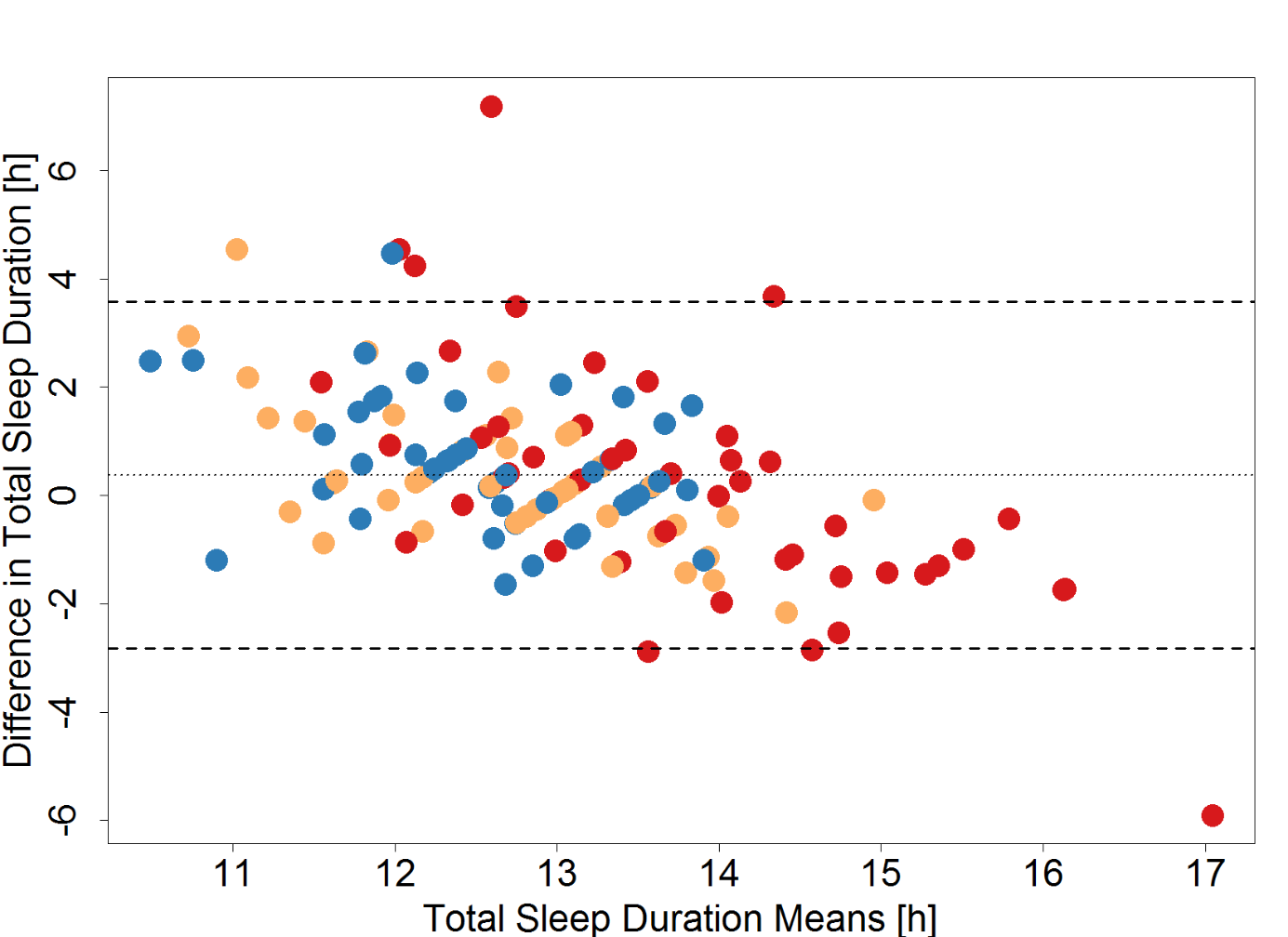

**Supplementary Figure 1.** Bland-Altman plots of *Total Sleep Duration* estimates from Sadeh algorithm and parental reports (Brief Infant sleep Questionnaire). Each infant is represented by three dots with color indicating age. While differences are ~ 0 h in the mean, the critical difference is 3.20 h, indicating large differences between parent reports and objectively measured sleep duration.

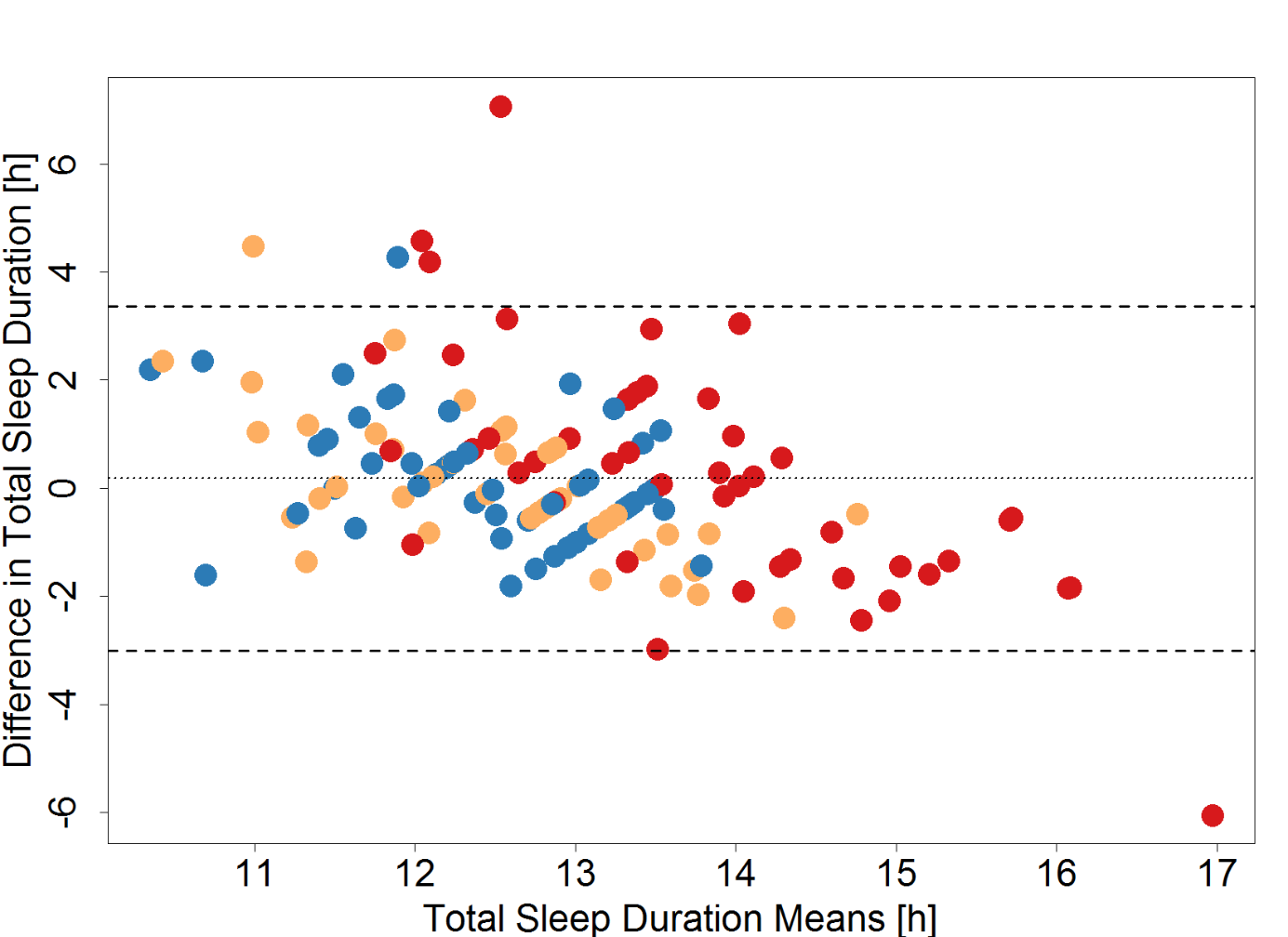

**Supplementary Figure 2.** Bland-Altman plots of *Total Sleep Duration* estimates from Oakley algorithm and parental reports (Brief Infant sleep Questionnaire). Each infant is represented by three dots with color indicating age. While differences are ~ 0 h in the mean, the critical difference is 3.19 h, indicating large differences between parent reports and objectively measured sleep duration.
